# A close-to-native structure of the synaptonemal complex

**DOI:** 10.1101/2021.05.17.444495

**Authors:** Rosario Ortiz, Olga M. Echeverría, Sergej Masich, Christer Höög, Abrahan Hernández-Hernández

## Abstract

Genetic variability in sexually reproducing organisms results from an exchange of genetic material between homologous chromosomes. The genetic exchange mechanism is dependent on the synaptonemal complex (SC), a protein structure localized between the homologous chromosomes. Current structural models of the SC are based on electron microscopy, super resolution, and expansion microscopy studies using chemical fixatives and sample dehydration of gonads, which are methodologies known to produce structural artifacts. We have developed a novel electron microscopy sample-preparation approach where pachytene cells are isolated from mouse testis by FACS, followed by cryo-fixation and cryo-substitution to achieve visualization of a close-to-native structure of the SC. We found that the central region of the SC was wider than previously recognized, and the transverse filaments more densely packed in the central region. Furthermore, we identified a structure nucleating the central element of the SC.

## Introduction

The Synaptonemal Complex (SC) is a conserved macromolecular proteinaceus scaffold that keeps homologous chromosomes (homologs) together until they have exchanged DNA during meiosis^1^. The SC is observed as a tripartite structure composed of two well-delimited lateral elements (LEs) and a central region (CR). Protruding out from the LEs, fine transverse filaments (TFs) cross the CR and, together with other SC-specific proteins, form the central element (CE) at a central position of the CR. Structural studies, as well as biochemical and molecular characterization of SC proteins in mice, *Drosophila*, *C. elegans* and budding yeast have led to several models for the structure and organization of the SC^2,3,12–21,4,22–31,5,32–34,6–11^.

Exchange of genetic material (meiotic recombination) depends on intact CR and CE structures as structural protein mutations result in recombination blockage, meiotic arrest and massive cell death of meiocytes^7,26,29,35–38^. There is still a dearth of detailed knowledge of the interplay between CR structure and meiotic recombination. Understanding the close-to-native structure of the CR may shed light on the recombination process and the role of the SC in this process. Structural studies aiming to delineate the fine structure of the SC and the relative localization of its components have been performed by electron microscopy (EM), superresolution and expansion microscopy^2,3,33,39,40,8,9,12,16,18,22,30,31^.

A common caveat with these structural studies has been the use of chemical fixatives and sample dehydration or sample spreading of the gonads prior to EM analysis or superresolution/expansion microscopy. These methods are known to produce artifacts in the fine structure of macromolecular complexes^41–44^. However, no attempts have been made to establish a cryo-fixation/substitution experimental setup followed by electron tomography in order to achieve close-to-native structure of the SC. Therefore, we developed a novel approach to study the close-to-native structure of the SC in mouse germ cells. In this procedure, we isolated and enriched cells in which the SC is fully formed (pachytene cells) by Fluorescence-activated Cell Sorting (FACS). We then implemented a High-Pressure Freezing and Freeze-Substitution protocol (HPF/FS) to enable cryo-fixation and cryo-substitution of these pachytene cells. Finally, we performed electron tomography (ET) of the SC in the cryo-preserved pachytene cells. We compared the structure of SCs in cryo-fixated pachytene cells versus SCs from the classical chemical fixation of seminiferous tubules from mice. We observed that the structure and organization of the SC components from cryo-fixated cells is different from its classical tripartite structure as reported so far. We found that the CR is wider and that the TFs seem to be more densely packed in the CR. Furthermore, we observed a central repetitive structure that nucleates the CE. Thus, our data provides (for the first time ever) a snapshot of the close-to-native structure of the SC.

## Results

### An experimental approach to elucidate the close-to-native structure of the synaptonemal complex

We established a workflow to describe the close-to-native structure of the SC in pachytene cells in mouse testis. FACS sorting of testicular cells and isolation of pachytene cells prior to HPF/FS (hence, FACS/HPS/FS) was used to remove constraints caused by limited freezing depth in tissues treated using the HPF technique. We compared the integrity of cellular structures in pachytene cells prepared using FACS/HPS/FS versus the classical chemical fixation methods applied to seminiferous tubules. It has been shown that a close-to-native structure of the Golgi apparatus and the nuclear pore complex can be visualized following HPF/FS preparation of cells^45–49^.We performed ultrathin sectioning of the pachytene cells and analyzed the integrity of the Golgi apparatus and the nuclear pore complex by EM in these cells. As was the case of studies of other cell types, we found the Golgi apparatus and nuclear pore complex to be better preserved in cryo-preserved pachytene cells than in chemically fixed seminiferous tubules (Fig. 1a-f).

**Fig. 1.**
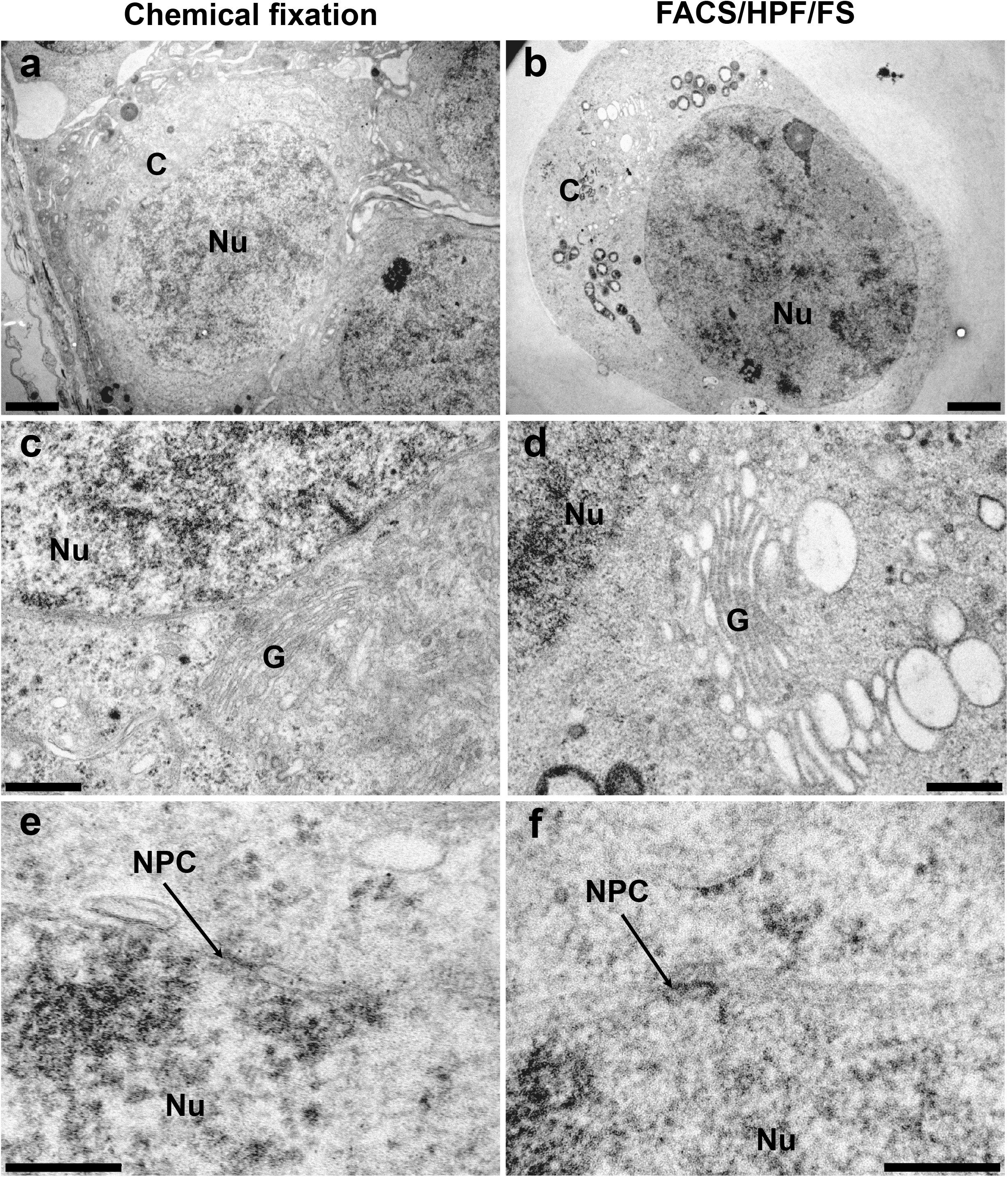
Cellular structures in chemically- and cryo-fixed samples. **a-b** Cytoplasm (C) and Nucleus (N) of pachytene cells. **c-d** Golgi apparatus in chemically-(Chemical fixation) or cryo-fixed (FACS/HPF/FS) samples, respectively. **e-f** Nuclear Pore Complex (NPC) in chemically- or cryo-fixed samples, respectively. Scale bars represent 2 μm in a and b, 500 nm in c and d, and 200 nm in e and f.

### The SC components in cryo-preserved pachytene cells are not as well-defined as in chemically fixed seminiferous tubules

The structure of the SC was first analyzed in chemically fixed seminiferous tubules and FACS-sorted pachytene cells. In a frontal view of the SC, we were able to clearly identify the borders of the LEs and the CE (Fig. 2a, c). These borders were more diffuse in SCs in cryo-preserved cells (Fig. 2b). The transverse filaments were also more distinct in pachytene cells in chemically fixed samples (Fig. 2a-c). Thus, it is likely that the sharply defined border for the different SC structures result from fixation artifacts following the use of chemicals.

**Fig. 2.**
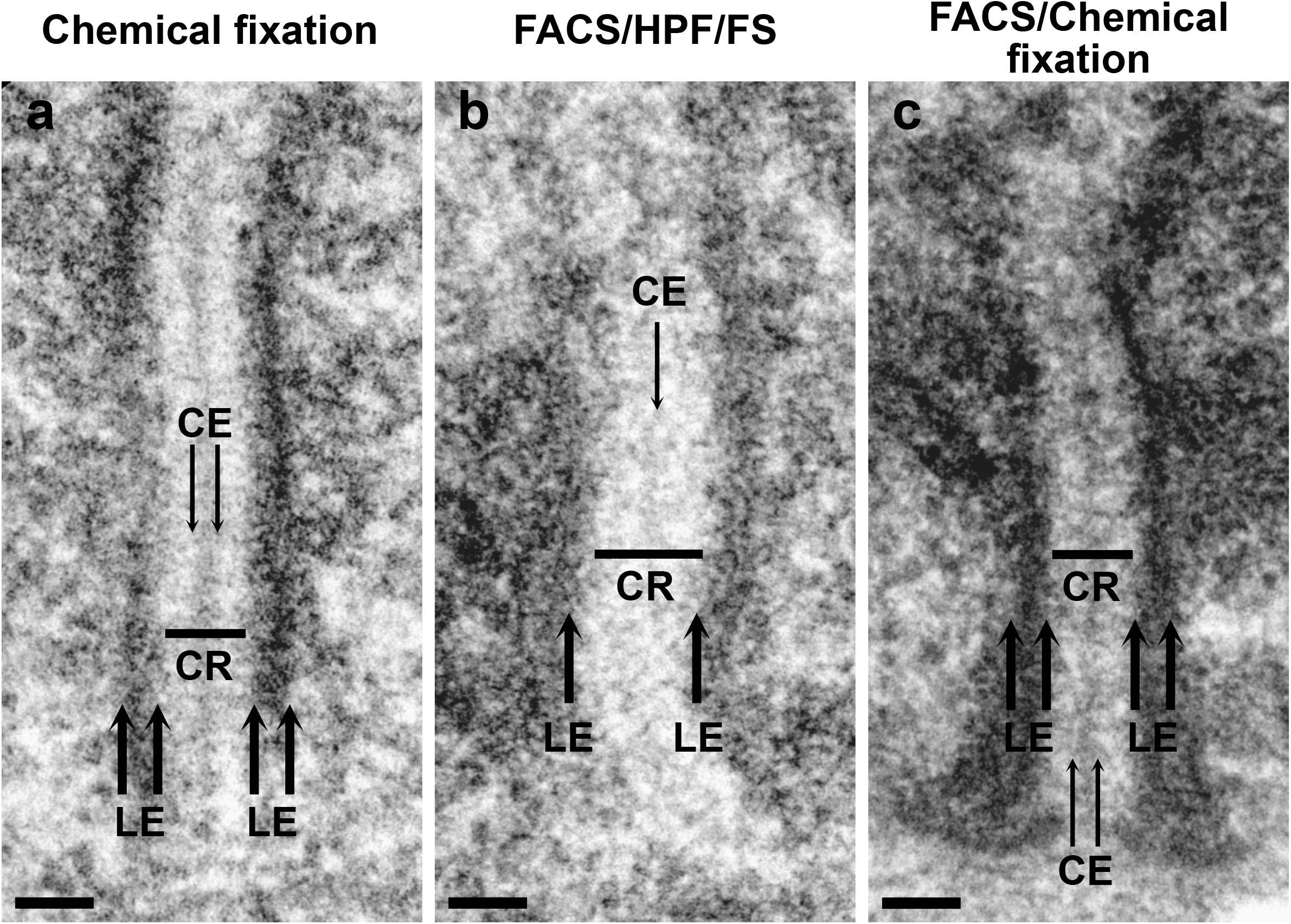
Structure of the SC in chemically- and cryo-fixed samples. **a** The SC’s tripartite organization with two lateral elements (LE) separated by a central region (CR) and in the middle the central element (CE), in pachytene cells after chemical fixation, dehydration and sectioning of seminiferous tubules. **b** The SC with LEs with an traceable border only at its inner side. The CR is clearly delimited whereas the CE is visible but without clear borders in pachytene cells after FACS isolation, high pressure freezing/freeze substitution (FACS/HPS/FS) and sectioning. **c** The SC displaying its classical tripartite organization in pachytene isolated by FACS and followed by chemical fixation and sectioning. Scale bar represents 100 nm.

We next performed double-tilt electron tomography of SCs in chemically fixed seminiferous tubules and in cryo-preserved pachytene cells. We manually segmented the SCs that were clearly defined in 4 and 5 tomograms from chemically fixed seminiferous tubules and cryo-preserved pachytene cells, respectively (Supplementary Videos 1-4 for chemically fixed samples and Supplementary Videos 5-9 for cryo-preserved samples). The outer and inner borders of the LEs and the borders of the CE in SCs from chemically fixed seminiferous tubules could be traced, whereas only the inner border of the LEs in SCs from cryo-preserved cells could be traced (Fig. 3a-d and Supplementary Fig. 1). Finally, we delineated TFs that fully cross the space between the inner side of the LE and the CE with a single line following their path (Fig. 3a-d and Supplementary Fig. 1). Thus, we found that the different sub-structures of the SC are less sharply outlined in cryo-preserved pachytene cells, representing a close-to native structure of the SC different than that resulting from the chemical fixation of meiotic cells.

**Fig. 3.**
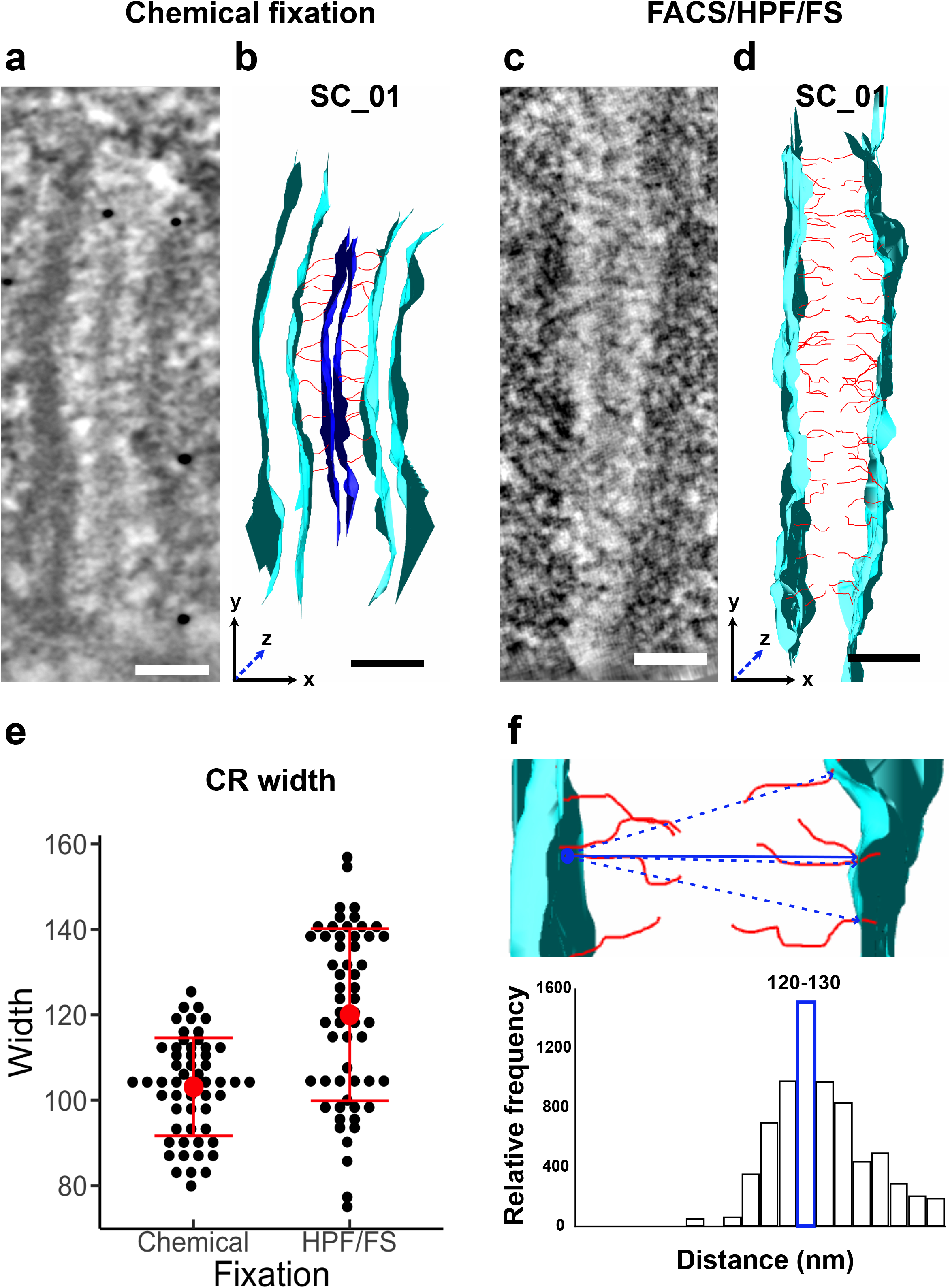
Electron tomography of the SC. **a-b** three-dimensional (3D) reconstruction (tomogram) and manual segmentation of a SC from seminiferous tubules with chemical fixation. The tomogram in panel a, has a depth along the Z axis of 33 nm. **c-d** three-dimensional (3D) reconstruction (tomogram) and manual segmentation of a SC from cryo-fixated pachytene cells (FACS/HPF/FS). The tomogram in panel c, has a depth along the Z axis of 36 nm. Scale bars represent 100 nm. **e** Dot plot displaying all the measurements of the CR width in samples with chemical fixation and FACS/HPS/FS. Red dots indicate the average and red lines indicate the standard deviation. **f** Top: neighbor density analysis (nda) computes the closest distance between LEs. Dotted blue lines represent the computed distances and the solid line the closest distance between one point in one LE against all the other points in the opposite LE. Bottom: bar chart of the preferred distance between LEs, the relative frequency of the computed distances is plotted on the “y” axis and the distance in bins of 10 nm on the “x” axis. The bin with most relative frequencies is highlighted in blue.

### The CR of the SC in cryo-preserved pachytene cells shows an extended width

We found the CR to be the only SC component to be clearly defined in cryo-preserved pachytene cells. We measured and compared CR width in different slices of the tomograms of frontally orientated SCs following the application of the two different fixative methods. The top, the middle and the bottom sections of the tomogram of SCs were selected for analysis (i.e. in the Z plane of the SC). Measurements of the width were taken every 100 nm along the Y axis of the SC (Supplementary Fig. 2). The width of the CR in cryo-preserved samples was estimated to be 120 nm on average, whereas the width of the CR of the SC in chemically fixed seminiferous tubules was estimated to be 103 nm on average (Fig. 3e). Thus, the width of the CR in SCs of cryo-preserved pachytene cells was approximately 16% wider.

### The organization of transverse filaments

Next, we analyzed the organization of the TFs of the SC in cryo-preserved pachytene cells and chemically fixed seminiferous tubules. A neighbor density analysis (nda) was used, which is a method used to analyze the spatial relationships between modeled objects in tomograms^50–52^. As proof of principle, to corroborate that the nda approach was suitable for our purposes, we challenged it by computing the preferred closest distance between LEs in cryo-preserved pachytene cells (Fig. 3f). We rationalized that the preferred closest distance between the LEs of the SC, defined by the contact points between the TFs and the inner border region of the LEs (Fig. 3f), should be similar to the average distance between the LEs obtained by direct measuring in the 3D structures. Indeed, the preferred closest distance between the LEs was positioned in the 120 to 130 nm range (Fig. 3f), similar to the 120 nm identified as the average distance between LEs (Fig. 3e). Next, the preferred closest distance between TFs in the 4 and 5 different 3D volumes of SCs from chemically fixed and cryo-preserved samples was analyzed. We found the preferred closest distance for TFs varied for SCs independently of the fixation protocol used (Fig. 4a-b). However, while the closest distances in tomograms for chemically fixed seminiferous tubules varied from 0-18 nm (Fig. 4a), the variation in cryo-fixed pachytene was only 0-8 nm (Fig. 4b), suggesting a higher density of TFs in SCs when analyzed by HPF/FS. Thus, cryo-preservation provides more information about TFs organization in the CR of the SC. Furthermore, the observation that there is not a constant preferred closest distance for the TFs in tomograms suggests that the disposition of the TFs in the CR is dynamic, as suggested by live cell imaging analysis of the SC in C. elegans^20^.

**Fig. 4.**
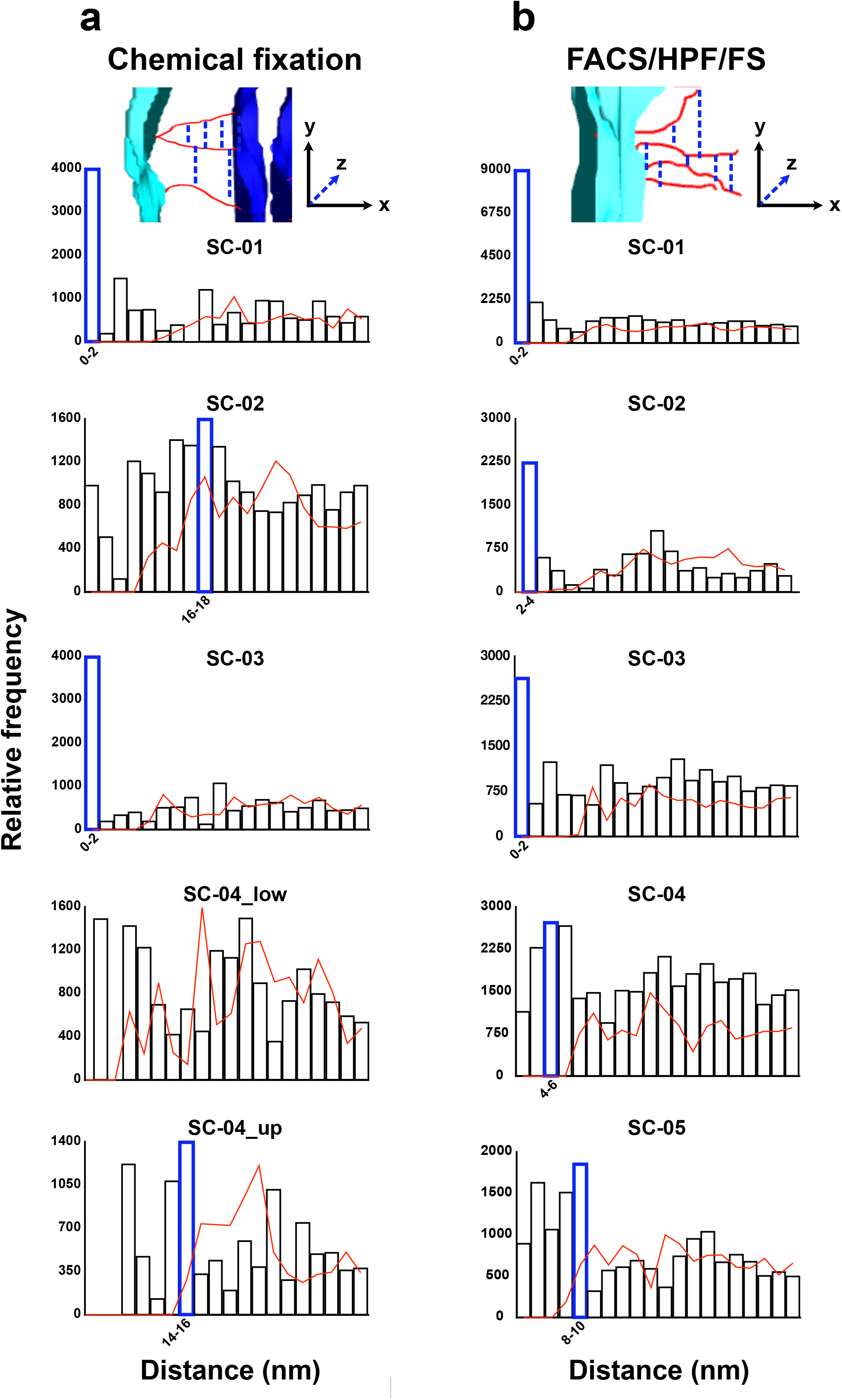
The preferred distance among TFs. **a** neighbor density analysis (nda) among TFs in SCs from seminiferous tubules with chemical fixation. Bar charts plotting the preferred distances in 4 tomograms are displayed. Tomogram 4 is split in low- and up-regions of the SC. **b** neighbor density analysis (nda) among TFs in SCs from cryo-preserved pachytene cells prepared by FACS/HPF/FS. Bar charts plotting the preferred distances in 5 tomograms are displayed. Preferred distance calculations when TFs are randomly shifted are displayed with red lines. The relative frequency of the computed distances is plotted on the “y” axis and the distance in bins of 2 nm on the “x” axis. The bins with most relative frequencies and that do not display random-like distribution are highlighted in blue.

Next, we performed sub-tomogram averaging^53,54^ of TFs from 4 tomograms from chemically fixed seminiferous tubules (55 and 62 single TFs placed to the left and right of the CE respectively, Supplementary Fig. 3a-c) and from 5 tomograms from cryo-preserved pachytene cells (103 and 94 single TFs placed to the left and right of the CE respectively, Supplementary Fig. 3d-f). We found no conserved organization for the averaged TFs using either fixation protocol (Fig 5. and Supplementary Videos 10-11), thus implying that individual TFs do not attain a stable conformation in the CR. Thus, our results support the view that the organization and TF structure in the CR of the SC is highly dynamic and/or flexible.

**Fig. 5.**
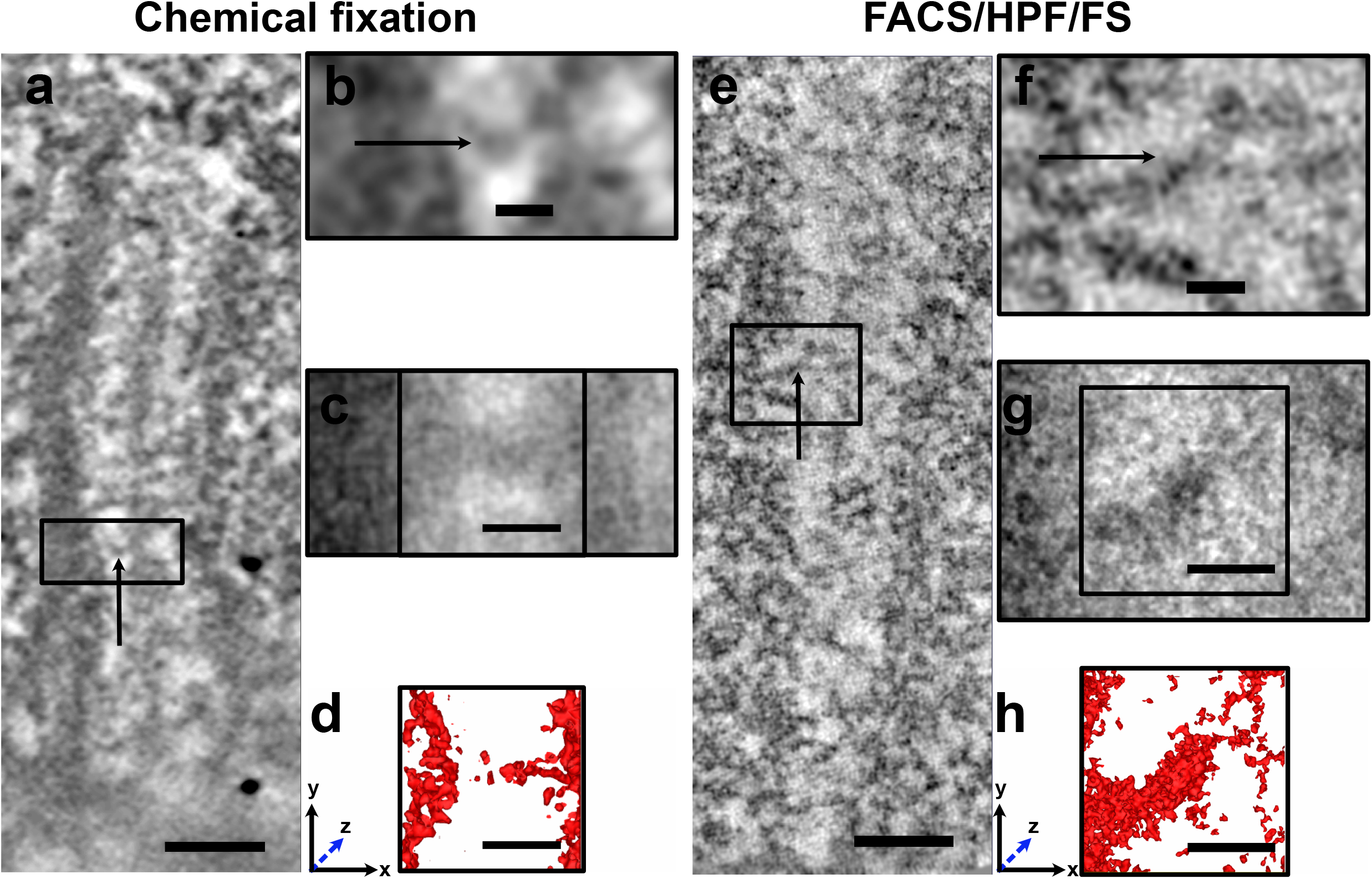
Sub-tomogram TF averaging. **a** one tomographic slide (0.78 nm) of a SC from seminiferous tubules with chemical fixation. The reference TF (arrow) used for sub-tomogram averaging is displayed in the rectangle. **b** Higher magnification of reference TF from panel a. **c** One tomographic slide (0.78 nm) of the structure obtained after averaging 55 TFs from seminiferous tubules with chemical fixation. **d** Automatic segmentation (surface rendering) of the averaged structure displayed in rectangle in d, with a depth of 10 nm along the “z” axis. **e** one tomographic slide (0.42 nm) of a SC from cryo-fixed pachytene cells (FACS/HPF/FS). The reference TF (arrow) used for sub-tomogram averaging is displayed in the rectangle. **f** Higher magnification of reference TF from panel e. **g** One tomographic slide (0.42 nm) of the structure obtained after averaging 103 TFs from cryo-fixed pachytene cells. **h** Automatic segmentation (surface rendering) of the averaged structure displayed in rectangle in g, with a depth of 10 nm along n the “z” axis. Scale bars represent 100 nm in a and c, and 20 nm in b-d, f-h.

### Analysis of the CE of the SC identified a central 10 nm structure

We observed a repeated structure across the CE of the CR of the SC in all tomograms from cryo-preserved pachytene cells, not apparent in chemically fixed seminiferous tubules (Fig. 6a-b). To further analyze this structure, we averaged 35 CE structures in cryo-preserved pachytene cells and identified an averaged structure that resembled the individual structures (Fig. 6c and Supplementary Video 12). The average structure does not reveal fine structural details; however, in every plane of the 3D structure, a round shape approximately 10 nm in width in the inner part of the CE is noticeable (Fig. 6d-g).

**Fig. 6.**
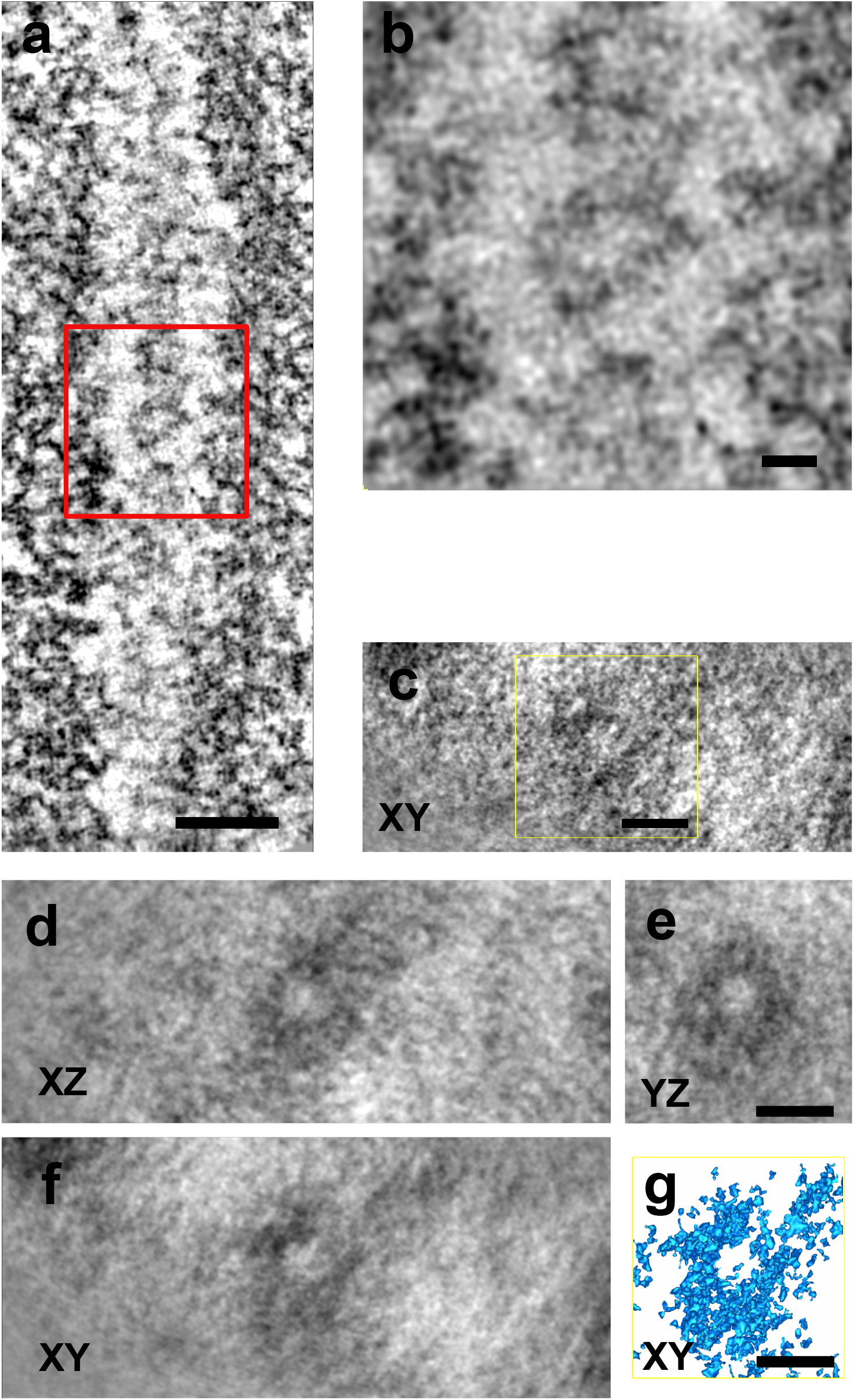
Sub-tomograms averaging of CE structures. **a** one tomographic slide (0.42 nm) of a SC from cryo-fixed pachytene cells (FACS/HPF/FS). The reference CE structure used for sub-tomogram averaging is displayed in the red rectangle. **b** Higher magnification of reference CE structure from panel a. **c** One tomographic slide (0.42 nm) of the structure obtained after averaging 35 CE structures from cryo-fixed pachytene cells. **d-f** Averaged CE structure on the XZ, YZ and XY axes of the tomogram with 8 nm of depth. **g** Automatic segmentation (surface rendering) of the averaged structure displayed in rectangle in c, with a depth of 8 nm along the “z” axis. Scale bars represent 100 nm in a, and 20 nm in b-g.

## Discussion

Cryo-fixation of the SC reveals a structural organization that differs in several ways from the SC structure resulting from the use of chemical fixation methods (Fig. 7). We show that the outer borders of the LEs and the CE are not well-defined, suggesting that the sharp definition of the borders in chemically fixed SCs are preparation artefacts. We found the distance between the inner part of the LEs and the CR to be wider than reported. The TFs seem to be more densely packed than previously recognized and an organized structure runs throughout its entire length of the CE (Fig 7).

**Fig. 7.**
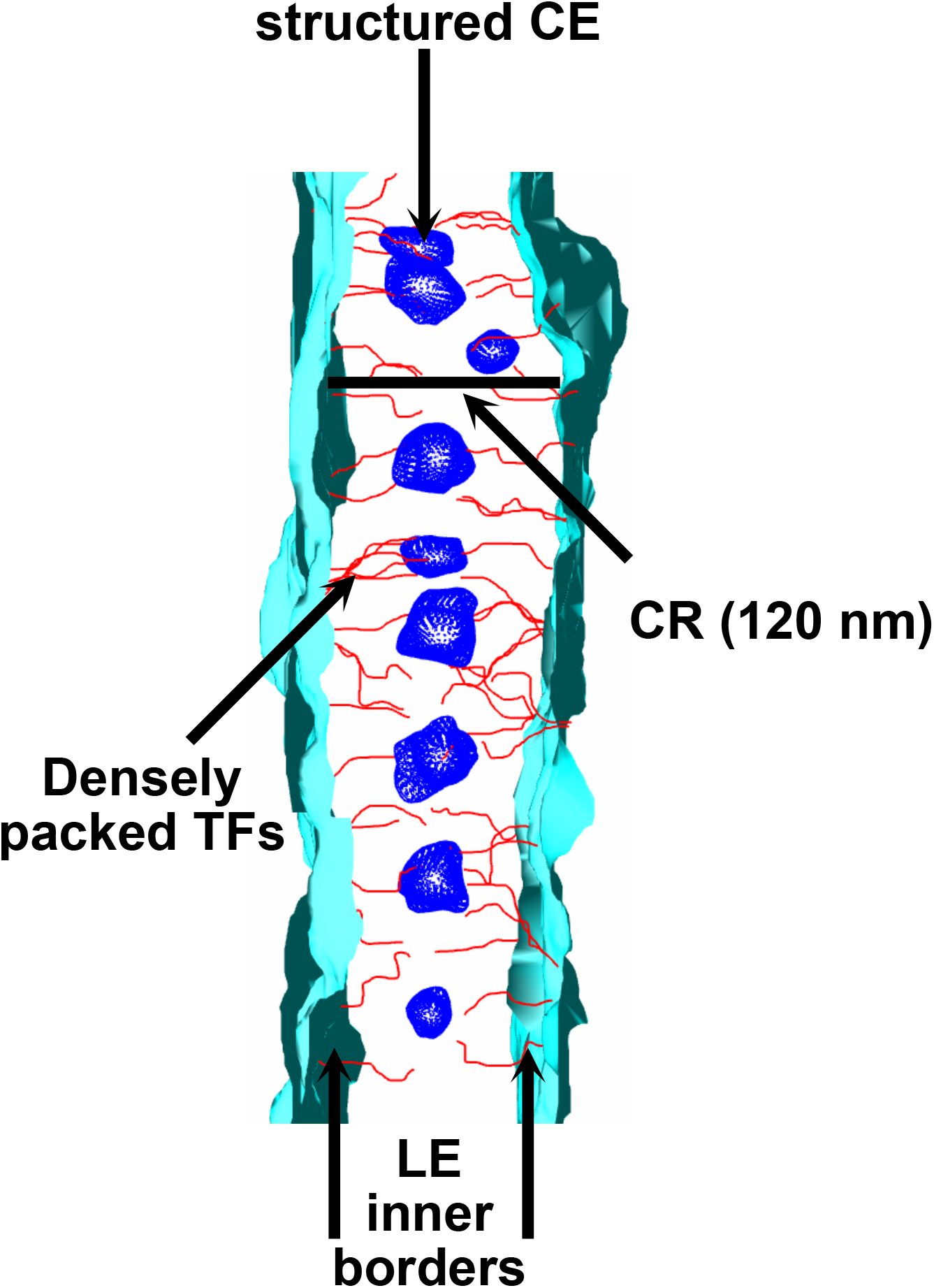
A snapshot of the close-to-native structure and organization of the SC.

Structural analysis of the SC by Immuno-EM, EM-tomography and expansion microscopy data^2,8,9,22^ using chemical fixatives is likely to introduce artifacts. These fixatives display a slow penetration speed relative to many cellular processes and may react chemically with cellular structures^42–44,55,56^. Thus, chemical fixatives used for conventional EM may result in the aggregation of proteins in a complex, such as the SC that is composed of multiple sub-structures merged together for an extended time period (the leptotene to pachytene stages of meiosis). Therefore, observation of a bi- or multilayered TFs organization may be a result of CR protein stacking due to an extended chemical fixation process. Additionally, preparation of SC spreads may also produce artifacts as the cells are subjected to swelling for several minutes prior to chemical fixation and spreading. In contrast, cryo-fixation of isolated pachytene cells takes place in 10-12 milliseconds, providing a snapshot of biological structures inside the cell^44,57^.

### A wider CR

It has been reported that the width of the CR of the SC is approximately 100 nm using chemical fixation methods of meiotic cells^25,31,39,58^. However, it has been suggested that the coiled-coiled region of the murine SYCP1 protein forming the TFs that span the distance between the LE and the CE is 90 nm long^13^ generating a theoretical CR width of 180 nm. The increased width of the CR found in cryo-fixed cells accommodates more of the proposed relative TF length.

### TF Organization

After sub-tomogram averaging, we did not observe any stable conformation of the average TFs. This is the expected scenario when averaging dynamic and/or flexible structures so that every targeted complex represents a unique conformational species^59^. Thus, dynamic TFs organization in the CR observed in this work is more comparable to the high mobility of SC components in live cell imaging studies. Theses TFs dynamics, in turn, may be dictated by the liquid-like crystal properties of the SC components^20^. However, we cannot completely rule out that sample staining in this close-to-native approach and/or low number of particles are making it difficult to visualize stable TF conformation in the CR, thereby precluding functional and physiological interpretations^59^. Thus, the development of methodologies allowing the visualization of the native TF conformation is required.

### CE organization

Contrary to the TFs, when we averaged the observed structures in the CE, we visualized a central structure with a doughnut shape of more or less 10 nm in diameter. This implies that CE proteins may be forming more stable structures that help to keep the LE of the SC together. CE stability may explain why some of these proteins can form filaments *in vitro*, resembling the CE, whereas the TF protein SYCP1 can only polymerize *in vivo* when overexpressed in the cytoplasm of cultured cells^8,60,61^.

Further methodological development that makes it possible to analyze the native SC structure should allow us to understand CR, TF, and CE structures as well as the LEs of the SC. Docking of atomic resolution crystallography maps of SYCP1 and CE proteins in the native structure of the SC should provide us with a high-resolution view of the SC. In addition, systematic 3D reconstructions of full-length SCs at different stages of maturation during meiosis will help us to understand how this structural scaffold promotes meiotic recombination.

## Methods

### Fluorescence Activated Cell Sorting (FACS) of Pachytene cells

Experimental procedures involving mice were approved by the Stockholm-North Animal Ethical Committee. Spermatogenic cell dissociation from testis, staining, gating and sorting of pachytene cells were done as reported above^62^.

### High-Pressure Freezing (HPF)

Sorted pachytene cells were pelleted at 3500 rpm at 4°C for 10 min and re-suspend in 600 μl of a 1:1 solution of modified HBSS (Hanks’ balanced salt solution, supplemented with 20 mM Hepes pH 7.2, 6.6 mM sodium pyruvate and 0.05% sodium lactate) and FBS (fetal bovine serum). Cells were pelleted once again and resuspended with 50 μl of 2% low-temperature gelling agarose at 37°C^63^ diluted in modified HBSS plus FBS (as mentioned above).

From this step on, solutions and plastic tips were kept in a heating block at 37°C. After centrifugation, 30 μl of the supernatant of agarose solution were removed and the pelleted cells were carefully mixed with the remaining agarose solution. The pachyte/agarose solution was briefly kept at 37°C until cryo-fixation. Ten μl of these pachytene cells in agarose were placed on a flat carrier (1.2 mm of diameter and 200 μm of depth) and high-pressure frozen in the EMPACT2 high-pressure freezer (in accordince with the manufacturer’s instructions, Leica Microsystems).

### Freeze-substitution (FS)

Frozen samples were freeze-substituted in the automatic Leica Microsystems EM AFS2. A freeze-substitution protocol that allows improved contrast of proteins and membranes was adapted for this work^46,47,64,65^ and carried out over a 7-day period as follows:

Frozen samples on the flat carriers were placed in a solution of acetone containing 0.5% tannic acid and 0.5% glutaraldehyde at −90°C for 48 hours. The temperature was gradually increased to −25°C in an 8-hour cycle and the samples were then rinsed three times with acetone and submerged in a solution of acetone with 2% osmium tetra-oxide plus 0.8% potassium ferrocyanide and 0.5% uranyl acetate for 4 hours. The temperature was then gradually increased to +4°C in a 2-hour cycle and the samples were rinsed three times with acetone. Samples were transferred to room temperature (RT) and infiltrated with a 1:1 mixture of propylene oxide/acetone for 5 min and 3 changes of propylene oxide for 30 min each. For embedding, the samples were submerged in a mixture of 1:1 propylene oxide/Durcupan overnight prior to incubation in freshly made Durcupan for 3 hours and finally, they were embedded and polymerized in Durcupan at 60°C for 72 hours.

### Chemical fixation of seminiferous tubules and sorted pachytene cells

Seminiferous tubules were processed as described above^8^. Briefly, small fragments of mice testes were fixed with 2.5% glutaraldehyde and 4% paraformaldehyde for 2 h at RT, post-fixed with 2% osmium tetroxide for 1 h, rinsed with cacodylate buffer, dehydrated through graded ethanol solutions, embedded in Durcupan and polymerized at 60°C for 72 hours.

Sorted pachytene cells were processed as mentioned above until they were pelleted and resuspended with 50 μl of 2% low-temperature gelling agarose at 37°C. Then chemical fixation with 2.5% glutaraldehyde and 4% paraformaldehyde was performed and further processing was done as for the seminiferous tubules.

### EM data acquisition, tomogram generation and analysis

For conventional TEM, 80 nm ultrathin sections were made using an ultra-microtome (Leica). Sections were placed on formvar/carbon-coated one-slot cupper grids (Agar Scientific), stained for 20 min in uranyl acetate and 10 min in lead citrate. Grids were observed with an electron microscope CM120 (FEI microscope) at 100 kV equipped with a side-entry MegaView III CCD camera to record the images and AnalySYS-3.2 software (Olympus Soft Imaging Solutions, Münster, Germany).

For electron tomography, 100nm-thick sections were placed on formvar/carbon-coated one-slot gold grids (Agar Scientific), previously incubated with 10 nm colloidal gold for alignment during tomogram reconstruction. The thickness of the sections was chosen accordingly to the thickness of the SC in the Z plane^22^. Grids were then post-stained with 3% uranyl acetate for 10 min. Initial screening to identify the SCs was performed with an electron microscope CM120 (FEI microscope) at 100 kV. For tomographic data collection, we used an FEG CM200 FEI microscope operating at 200 kV. The microscope was equipped with a cooled slow scan 2048 x 2048 TemCam F214 CCD camera and EM-MENU software for automated data collection (TVIPS, Gauting, Germany). The sections were pre-irradiated for 20 min before data collection at the beam intensity corresponding to the actual data acquisition. 260 images (double-tilt from −65° to +65° with one-degree increments) were collected and used for alignment and back projection reconstruction with IMOD software^66,67^. Signal-to-noise ratio in the tomograms was improved by SIRT iterative reconstruction in IMOD. Pixel sizes for the tomograms were 0.7838 nm at 23,000 magnification and 0.3248 nm at 50,000 magnification, for chemically- and cryo-fixed samples, respectively. Manual segmentation was done with IMOD drawing tools and the neighbor density analysis was taken and modified from previous reports^51,52^. TF and CE structure averaging was done according to the PEET (Particle Estimation for Electron Tomography) analysis tutorials^53,54^.

## Acknowledgements

We would like to thank the Electron Tomography Facility at Karolinska Institutet for providing its services. This work was supported by grants from the Swedish Research Council (2017-01853 to C.H) and the Hospital Infantil de México Federico Gómez (HIM/2016/096 SSA 1289 and HIM/2018/079 SSA 1518 to A.H.H).

## Author Contributions

A.H.H and C.H conceived the experiments. A.H.H designed and performed most of the experiments. R.O performed sample sectioning and data analysis.

S.M performed data acquisition and tomogram generation. C.H, O.M.E, and A.H.H contributed with reagents, equipment and funding. A.H.H and C.H supervised the experiments and data analyses. AHH and C.H wrote the manuscript. All the authors discussed the data and reviewed the manuscript.

## References

1. Zickler, D. & Kleckner, N. Recombination, pairing, and synapsis of homologs during meiosis. Cold Spring Harb. Perspect. Biol. 7, 1–28 (2015).

2. Schücker, K., Holm, T., Franke, C., Sauer, M. & Benavente, R. Elucidation of synaptonemal complex organization by super-resolution imaging with isotropic resolution. Proc. Natl. Acad. Sci. U. S. A. 112, 2029–2033 (2015).

3. Xu, H. et al. Molecular organization of mammalian meiotic chromosome axis revealed by expansion STORM microscopy. Proc. Natl. Acad. Sci. U. S. A. 116, 18423–18428 (2019).

4. Syrjänen, J. L., Pellegrini, L. & Davies, O. R. A molecular model for the role of SYCP3 in meiotic chromosome organisation. Elife 3, (2014).

5. Biswas, U., Hempel, K., Llano, E., Pendas, A. & Jessberger, R. Distinct Roles of Meiosis-Specific Cohesin Complexes in Mammalian Spermatogenesis. PLoS Genet. 12, e1006389 (2016).

6. Agostinho, A. & Höög, C. REC8 density along chromosomes prevents illegitimate synapsis. Cell cycle (Georgetown, Tex.) vol. 15 2543–2544 (2016).

7. Gómez-H, L. et al. C14ORF39/SIX6OS1 is a constituent of the synaptonemal complex and is essential for mouse fertility. Nat. Commun. 7, 13298 (2016).

8. Hernández-Hernández, A. et al. The central element of the synaptonemal complex in mice is organized as a bilayered junction structure. J. Cell Sci. 129, 2239–2249 (2016).

9. Spindler, M.-C., Filbeck, S., Stigloher, C. & Benavente, R. Quantitative basis of meiotic chromosome synapsis analyzed by electron tomography. Sci. Rep. 9, 16102 (2019).

10. Bolcun-Filas, E. & Schimenti, J. C. Genetics of meiosis and recombination in mice. Int. Rev. Cell Mol. Biol. 298, 179–227 (2012).

11. Cahoon, C. K. & Hawley, R. S. Regulating the construction and demolition of the synaptonemal complex. Nat. Struct. Mol. Biol. 23, 369–377 (2016).

12. Cahoon, C. K. et al. Superresolution expansion microscopy reveals the three-dimensional organization of the Drosophila synaptonemal complex. Proc. Natl. Acad. Sci. U. S. A. 114, E6857–E6866 (2017).

13. Dunce, J. M. et al. Structural basis of meiotic chromosome synapsis through SYCP1 self-assembly. Nat. Struct. Mol. Biol. 25, 557–569 (2018).

14. Dunne, O. M. & Davies, O. R. Molecular structure of human synaptonemal complex protein SYCE1. Chromosoma 128, 223–236 (2019).

15. Hurlock, M. E. et al. Identification of novel synaptonemal complex components in C. elegans. J. Cell Biol. 219, (2020).

16. Köhler, S., Wojcik, M., Xu, K. & Dernburg, A. F. Superresolution microscopy reveals the three-dimensional organization of meiotic chromosome axes in intact Caenorhabditis elegans tissue. Proc. Natl. Acad. Sci. U. S. A. 114, E4734–E4743 (2017).

17. Rong, M., Matsuda, A., Hiraoka, Y. & Lee, J. Meiotic cohesin subunits RAD21L and REC8 are positioned at distinct regions between lateral elements and transverse filaments in the synaptonemal complex of mouse spermatocytes. J. Reprod. Dev. 62, 623–630 (2016).

18. Schücker, K., Sauer, M. & Benavente, R. Superresolution imaging of the synaptonemal complex. Methods Cell Biol. 145, 335–346 (2018).

19. Wang, Y. et al. Combined expansion microscopy with structured illumination microscopy for analyzing protein complexes. Nat. Protoc. 13, 1869–1895 (2018).

20. Rog, O., Köhler, S. & Dernburg, A. F. The synaptonemal complex has liquid crystalline properties and spatially regulates meiotic recombination factors. Elife 6, (2017).

21. Fraune, J. et al. Evolutionary history of the mammalian synaptonemal complex. Chromosoma 125, 355–360 (2016).

22. Zwettler, F. U. et al. Tracking down the molecular architecture of the synaptonemal complex by expansion microscopy. Nat. Commun. 11, 3222 (2020).

23. Gao, J. & Colaiacovo, M. P. Zipping and Unzipping: Protein Modifications Regulating Synaptonemal Complex Dynamics. Trends Genet 34, 232–245 (2018).

24. Pyatnitskaya, A., Borde, V. & De Muyt, A. Crossing and zipping: molecular duties of the ZMM proteins in meiosis. Chromosoma (2019) doi:10.1007/s00412-019-00714-8.

25. Westergaard, M. & von Wettstein, D. The synaptinemal complex. Annu Rev Genet 6, 71–110 (1972).

26. Bolcun-Filas, E. et al. SYCE2 is required for synaptonemal complex assembly, double strand break repair, and homologous recombination. J Cell Biol 176, 741–747 (2007).

27. Fraune, J., Schramm, S., Alsheimer, M. & Benavente, R. The mammalian synaptonemal complex: protein components, assembly and role in meiotic recombination. Exp Cell Res 318, 1340–1346 (2012).

28. Hamer, G. et al. Characterization of a novel meiosis-specific protein within the central element of the synaptonemal complex. J Cell Sci 119, 4025–4032 (2006).

29. Bolcun-Filas, E. et al. Mutation of the mouse Syce1 gene disrupts synapsis and suggests a link between synaptonemal complex structural components and DNA repair. PLoS Genet 5, e1000393 (2009).

30. Schmekel, K., Skoglund, U. & Daneholt, B. The three-dimensional structure of the central region in a synaptonemal complex: a comparison between rat and two insect species, Drosophila melanogaster and Blaps cribrosa. Chromosoma 102, 682–692 (1993).

31. Ortiz, R., Kouznetsova, A., Echeverria-Martinez, O. M., Vazquez-Nin, G. H. & Hernandez-Hernandez, A. The width of the lateral element of the synaptonemal complex is determined by a multilayered organization of its components. Exp Cell Res 344, 22–29 (2016).

32. Solari, A. J. & Moses, M. J. The structure of the central region in the synaptonemal complexes of hamster and cricket spermatocytes. J Cell Biol 56, 145–152 (1973).

33. Collins, K. A. et al. Corolla is a novel protein that contributes to the architecture of the synaptonemal complex of Drosophila. Genetics 198, 219–228 (2014).

34. Humphryes, N. et al. The Ecm11-Gmc2 complex promotes synaptonemal complex formation through assembly of transverse filaments in budding yeast. PLoS Genet 9, e1003194 (2013).

35. de Vries, F. A. et al. Mouse Sycp1 functions in synaptonemal complex assembly, meiotic recombination, and XY body formation. Genes Dev 19, 1376–1389 (2005).

36. Geisinger, A. & Benavente, R. Mutations in Genes Coding for Synaptonemal Complex Proteins and Their Impact on Human Fertility. Cytogenet Genome Res 150, 77–85 (2016).

37. Hamer, G. et al. Progression of meiotic recombination requires structural maturation of the central element of the synaptonemal complex. J Cell Sci 121, 2445–2451 (2008).

38. Schramm, S. et al. A novel mouse synaptonemal complex protein is essential for loading of central element proteins, recombination, and fertility. PLoS Genet 7, e1002088 (2011).

39. Schmekel, K. & Daneholt, B. The central region of the synaptonemal complex revealed in three dimensions. Trends Cell Biol 5, 239–242 (1995).

40. Schmekel, K., Wahrman, J., Skoglund, U. & Daneholt, B. The central region of the synaptonemal complex in Blaps cribrosa studied by electron microscope tomography. Chromosoma 102, 669–681 (1993).

41. Schmekel, K. et al. Organization of SCP1 protein molecules within synaptonemal complexes of the rat. Exp Cell Res 226, 20–30 (1996).

42. Shiurba, R. Freeze-substitution: origins and applications. Int. Rev. Cytol. 206, 45–96 (2001).

43. Studer, D., Humbel, B. M. & Chiquet, M. Electron microscopy of high pressure frozen samples: bridging the gap between cellular ultrastructure and atomic resolution. Histochem Cell Biol 130, 877–889 (2008).

44. Vanhecke, D., Graber, W. & Studer, D. Close-to-native ultrastructural preservation by high pressure freezing. Methods Cell Biol 88, 151–164 (2008).

45. Fiserova, J., Spink, M., Richards, S. A., Saunter, C. & Goldberg, M. W. Entry into the nuclear pore complex is controlled by a cytoplasmic exclusion zone containing dynamic GLFG-repeat nucleoporin domains. J. Cell Sci. 127, 124–136 (2014).

46. Giddings, T. H. Freeze-substitution protocols for improved visualization of membranes in high-pressure frozen samples. J. Microsc. 212, 53–61 (2003).

47. Jiménez, N. et al. Tannic acid-mediated osmium impregnation after freeze-substitution: a strategy to enhance membrane contrast for electron tomography. J. Struct. Biol. 166, 103–106 (2009).

48. Marsh, B. J. & Pavelka, M. Viewing Golgi structure and function from a different perspective--insights from electron tomography. Methods Cell Biol. 118, 259–279 (2013).

49. McDonald, K. L. Out with the old and in with the new: rapid specimen preparation procedures for electron microscopy of sectioned biological material. Protoplasma 251, 429–448 (2014).

50. Hata, S. et al. The balance between KIFC3 and EG5 tetrameric kinesins controls the onset of mitotic spindle assembly. Nat. Cell Biol. 21, 1138–1151 (2019).

51. Marsh, B. J., Mastronarde, D. N., Buttle, K. F., Howell, K. E. & McIntosh, J. R. Organellar relationships in the Golgi region of the pancreatic beta cell line, HIT-T15, visualized by high resolution electron tomography. Proc. Natl. Acad. Sci. U. S. A. 98, 2399–2406 (2001).

52. Höög, J. L. et al. Organization of interphase microtubules in fission yeast analyzed by electron tomography. Dev. Cell 12, 349–361 (2007).

53. Heumann, J. M., Hoenger, A. & Mastronarde, D. N. Clustering and variance maps for cryo-electron tomography using wedge-masked differences. J. Struct. Biol. 175, 288–299 (2011).

54. Nicastro, D. et al. The molecular architecture of axonemes revealed by cryoelectron tomography. Science 313, 944–948 (2006).

55. Li, Y. et al. The effects of chemical fixation on the cellular nanostructure. Exp. Cell Res. 358, 253–259 (2017).

56. Dempster, W. T. Rates of penetration of fixing fluids. Am. J. Anat. 107, 59–72 (1960).

57. McEwen, B. F., Renken, C., Marko, M. & Mannella, C. Chapter 6: Principles and practice in electron tomography. Methods Cell Biol. 89, 129–168 (2008).

58. Moses, M. J. SYNAPTINEMAL COMPLEX. Annu. Rev. Genet. 2, 363–412 (1968).

59. Basanta, B., Chowdhury, S., Lander, G. C. & Grotjahn, D. A. A guided approach for subtomogram averaging of challenging macromolecular assemblies. J. Struct. Biol. X 4, 100041 (2020).

60. Davies, O. R., Maman, J. D. & Pellegrini, L. Structural analysis of the human SYCE2-TEX12 complex provides molecular insights into synaptonemal complex assembly. Open Biol 2, 120099 (2012).

61. Ollinger, R., Alsheimer, M. & Benavente, R. Mammalian protein SCP1 forms synaptonemal complex-like structures in the absence of meiotic chromosomes. Mol Biol Cell 16, 212–217 (2005).

62. Bastos, H. et al. Flow cytometric characterization of viable meiotic and postmeiotic cells by Hoechst 33342 in mouse spermatogenesis. Cytom. A 65, 40–49 (2005).

63. Sobol, M. A., Philimonenko, V. V, Philimonenko, A. A. & Hozak, P. Quantitative evaluation of freeze-substitution effects on preservation of nuclear antigens during preparation of biological samples for immunoelectron microscopy. Histochem Cell Biol 138, 167–177 (2012).

64. Cotta-Pereira, G., Rodrigo, F. G. & David-Ferreira, J. F. The use of tannic acid-glutaraldehyde in the study of elastic and elastic-related fibers. Stain Technol. 51, 7–11 (1976).

65. Bonilla, E. Staining of transverse tubular system of skeletal muscle by tannic acid-glutaraldehyde fixation. J. Ultrastruct. Res. 162–165 (1977) doi:10.1016/s0022-5320(77)90028-4.

66. Kremer, J. R., Mastronarde, D. N. & McIntosh, J. R. Computer visualization of three-dimensional image data using IMOD. J Struct Biol 116, 71–76 (1996).

67. Mastronarde, D. N. Dual-axis tomography: an approach with alignment methods that preserve resolution. J Struct Biol 120, 343–352 (1997).

