## Supplementary Figures Legends for "A close-to-native structure of the synaptonemal complex"

**Supplementary figure legends**

**Supplementary Fig 1. Manual segmentation of SCs.** **a-c** Manual segmentation of SCs from seminiferous tubules with chemical fixation. **d-g** Manual segmentation of SCs from cryo-fixated pachytene cells (FACS/HPF/FS). Scale bars represent 100 nm

**Supplementary Fig 2. Measurements of CR width in tomograms of frontally oriented SCs.** The top, middle and bottom sections of the tomogram of SCs were selected for analysis (i.e., in the Z plane of the SC). The inner borders of the LEs are delineated with magenta lines. Measurements of the width were taken every 100 nm along the Y axis of the SC (as displayed by red lines). **Top:** sections of tomograms of SCs from seminiferous tubules with chemical fixation. **Bottom:** sections of tomograms of SCs from cryo-fixated pachytene cells. Scale bars represent 100 nm

**Supplementary Fig 3. TFs used for sub-tomogram averaging. a** one tomographic slide (0.78 nm) of a SC from seminiferous tubules with chemical fixation. The reference TF used for sub-tomogram averaging is displayed in the rectangle. **b** Red lines in the tomogram display the TFs that fully cross the space between the LE and the CE and that were used for sub-tomogram averaging of TFs to the left of the CE. **c** one tomographic slide (0.78 nm) of a SC from seminiferous tubules with chemical fixation. The reference TF used for sub-tomogram averaging is displayed in the rectangle. Red lines in tomogram display the TFs that fully cross the space between the LE and the CE used for TF sub-tomogram averaging to the right of the CE. **d** One tomographic slide (0.42 nm) of a SC from cryo-fixed pachytene cells (FACS/HPF/FS). The reference TFs used for sub-tomogram averaging to the left and right sides of the CE are displayed in the red and blue rectangle, respectively. **e-f** Red lines in the tomogram display the TFs that fully cross the space between the LE and the CE and that were used for sub-tomogram averaging of TFs to the left and to the right of the CE, respectively. Scale bars represent 100 nm

**Supplementary videos 1-4.** Tomograms with manual segmentation for 4 SCs from chemically fixed seminiferous tubules. Scale bars represent 100 nm

**Supplementary videos 5-9.** Tomograms with manual segmentation for 5 SCs from cryo-preserved samples. Scale bars represent 100 nm

**Supplementary video 10.** Sub-tomogram averaging of TFs from chemically fixed seminiferous tubules. Scale bar represents 20 nm

**Supplementary video 11.** Sub-tomogram averaging of TFs from cryo-preserved samples. Scale bar represents 20 nm

**Supplementary video 12.** Sub-tomogram averaging of structure found in the CE from cryo-preserved samples. Scale bar represents 20 nm
