## Supplementary figures and images for "A close-to-native structure of the synaptonemal complex"

### Supplementary Figure 1

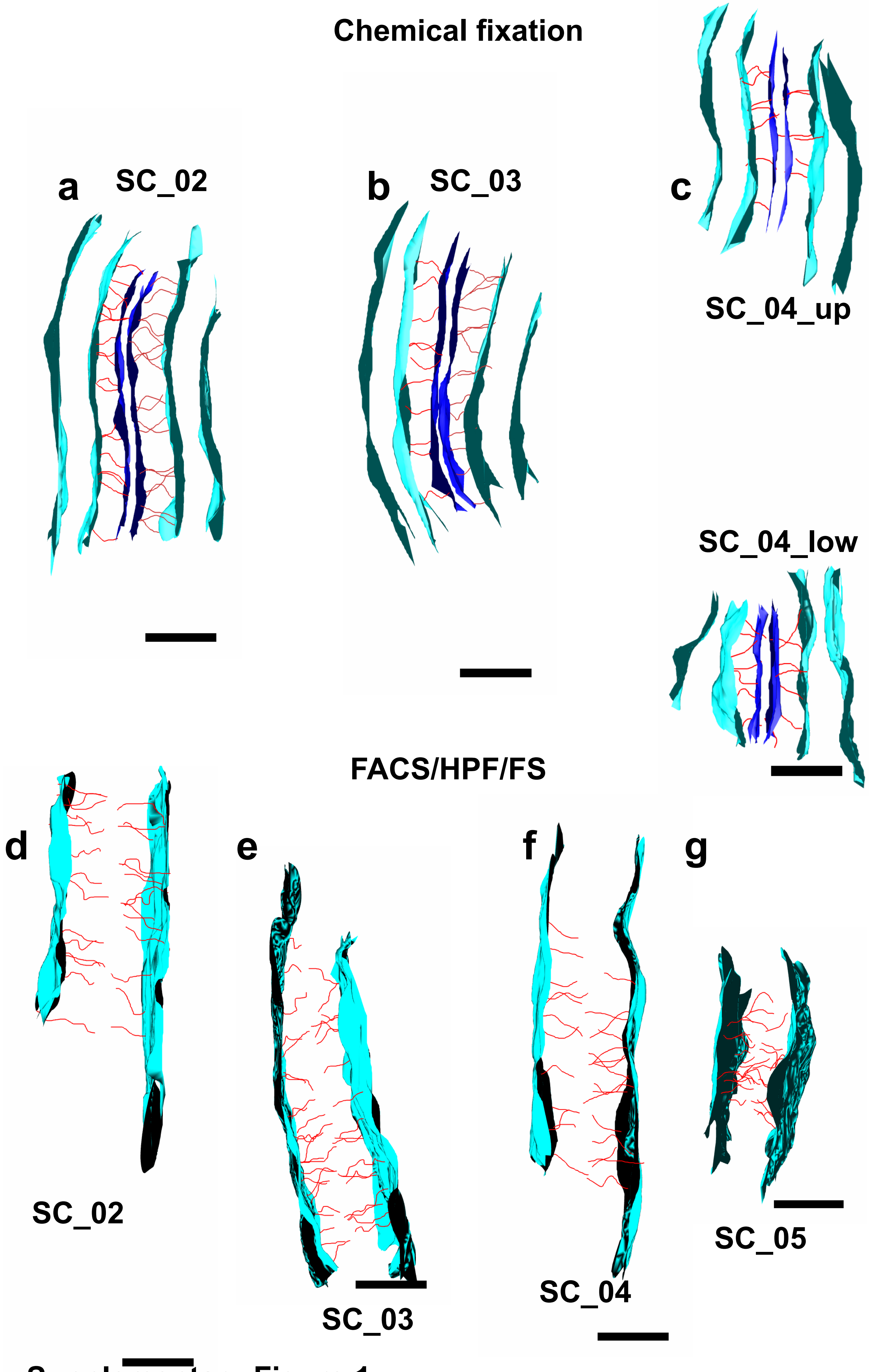


Supplementary Figure 1

### Supplementary Figure 2

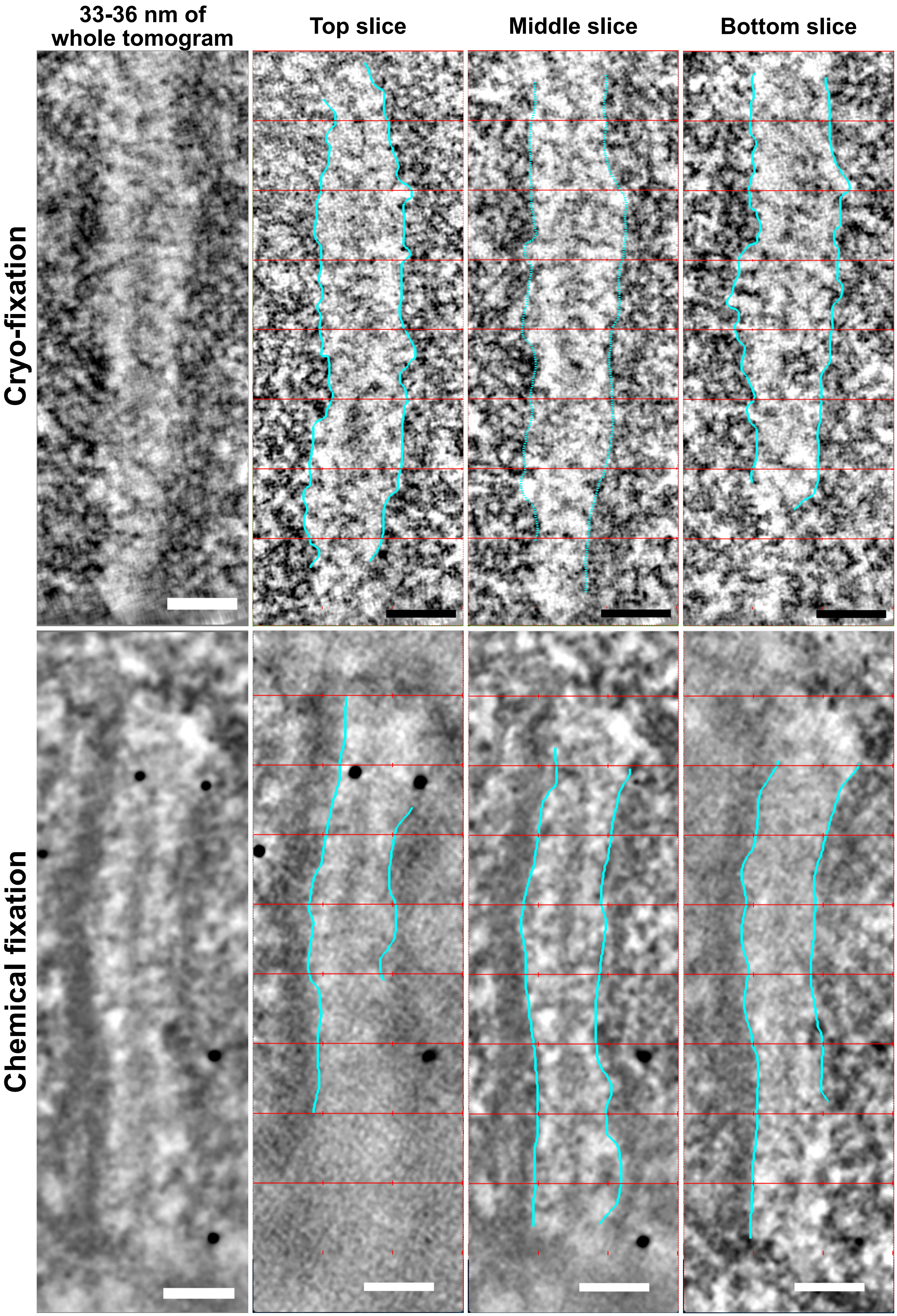


Supplementary Fig 2

### Supplementary Figure 3

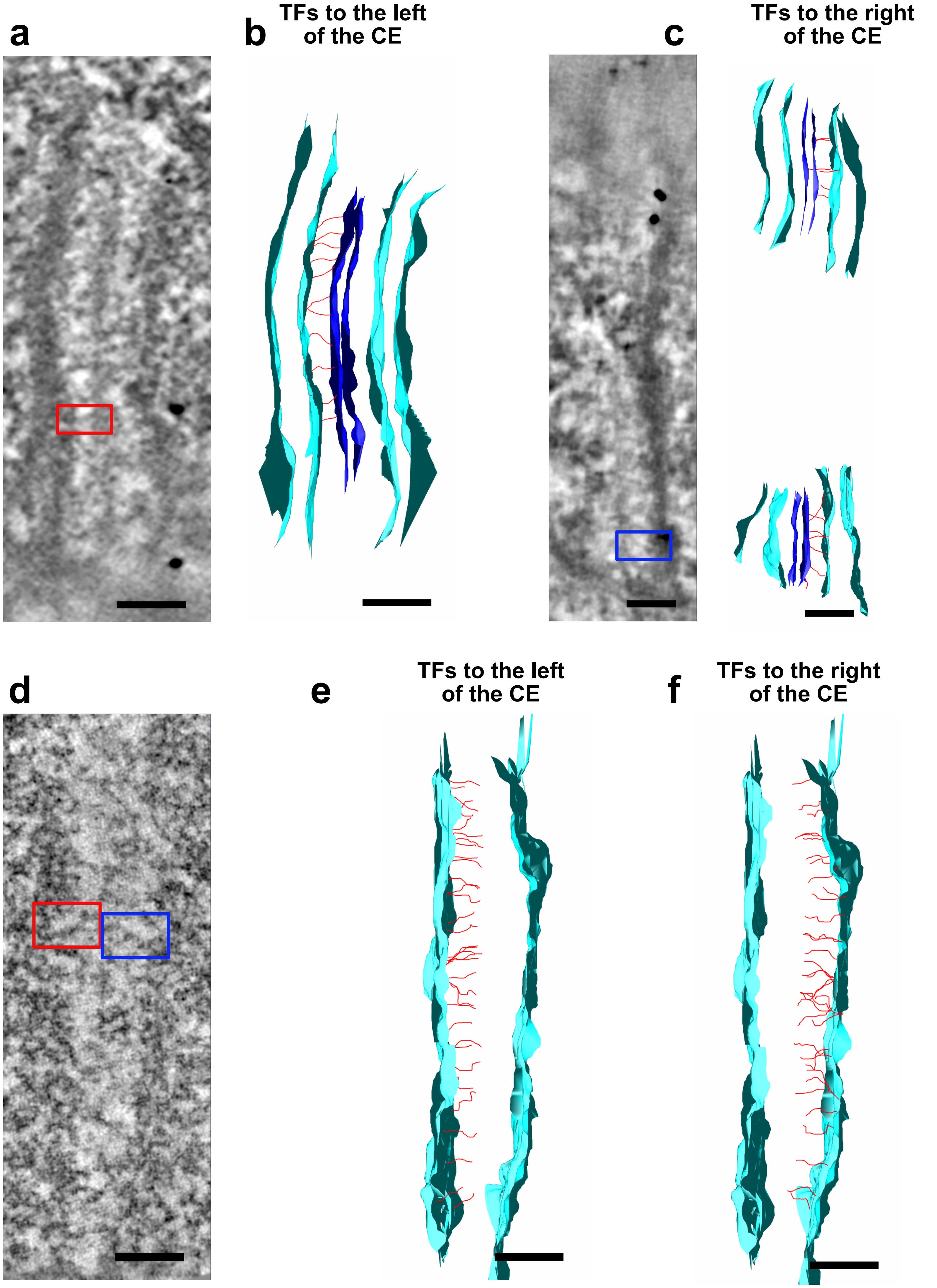


Supplementary Figure 3
